# Embedded Enzyme Nanoclusters Depolymerize Polyesters via Chain-End Mediated Processive Degradation

**DOI:** 10.1101/2020.04.25.052050

**Authors:** Christopher DelRe, Junpyo Kwon, Philjun Kang, Le Ma, Aaron Hall, Zhiyuan Ruan, Kyle Zolkin, Tim Li, Robert O. Ritchie, Ting Xu

## Abstract

Many bioactive elements, long perceived as non-viable for material development, are now emerging as viable building blocks to encode material lifecycle and to ensure our harmonious existence with nature. Yet, there is a significant knowledge gap on how bio-elements interface with synthetic counterparts and function outside of their native environments. Here, we show that when enzymes are dispersed as nanoclusters confined within macromolecular matrices, their reaction kinetics, pathway, and substrate selectivity can be modulated to achieve programmable polymer degradation down to repolymerizable small molecules. Specifically, when enzyme nanoclusters are dispersed in trace amount (~0.02 *wt*%) in polyesters, i.e. poly(caprolactone) (PCL) and poly(lactic acid) (PLA), chain-end mediated processive depolymerization can be realized, leading to scalable bioactive plastics for efficient sorting, such as recovery of precious metal filler from flexible electronics. Present studies demonstrate that when the enzyme is confined at dimensions similar to that of polymer chains, their behaviors are governed by the polymer conformation, segmental dynamic and thermal history, highlighting the importance to consider bioactive plastics differently from solution enzymology.

---

When it comes to ecological harmony, we envy nature’s ability to program and coordinate numerous complex processes to achieve system-wide, long-term sustainability (*1–4*). Synthetic biology opened a viable path to repurpose biomachineries toward new material design. Embedding enzymes and/or biomachineries capable of reacting with plastics (*5–7*) can afford on-demand backbone or side-chain modification and/or programmable degradation to enhance plastics’ environmental compatibility during manufacture/utilization (*8–10*). However, the potential of bioactive plastics is limited by the knowledge gap in biocatalysis in non-native environments (*11, 12*) and, in particular, achieving precise reaction control when the substrates are macromolecules acting as the host matrix.

The enzymatic activity depends on the protein structure, substrate binding, and reactivity at the active site (*13*) as schematically shown in **Fig. 1**. When a piece of solid plastic is immersed in an enzyme solution, such as in landfills, enzymes have sufficient translational diffusivity and conformational flexibility to bind to and catalyze reactions on the exposed polymer segments (*14, 15*) (**Fig. 1a**). However, the rate of surface erosion is so slow that even so called biodegradable plastics, such as poly(lactic acid) (PLA), are essentially non-degradable (*4, 16*). Accelerated random chain scissions have been achieved when the enzyme was dispersed as large enzyme/surfactant clusters via reverse micelles; however, the reaction proceeds preferentially in the amorphous domain (*17*), resulting in enzyme leaching and the formation of highly crystalline microplastic particles (*9*). When enzymes are confined at the length scales comparable to polymer chain dimensions, the kinetics and pathways of enzymatic reaction may vary (**Fig. 1b**). Thermodynamically, depolymerization of semi-crystalline polymers in solid state has several unique beneficial factors. The multi-pair inter-segment interactions in the crystalline domain provide a versatile handle to modulate chain end availability. Crystallized polymers have much lower conformational entropy and thus, a higher driving force for depolymerization than those in an amorphous state. Kinetically, substrate binding rather than the reaction rate may become the rate limiting step (**Fig. 1c**) due to reduced mobilities of both confined enzyme (*10–12, 18*) and semicrystalline polymer (*19, 20*). For semicrystalline polymers, which represent the majority of plastic production (*21*), the chain ends residing in the amorphous domain can be designed as the preferred binding sites such that processive depolymerization rather than random chain scission can be realized, leading to complete polymer degradation with defined byproducts.

**Fig. 1.**
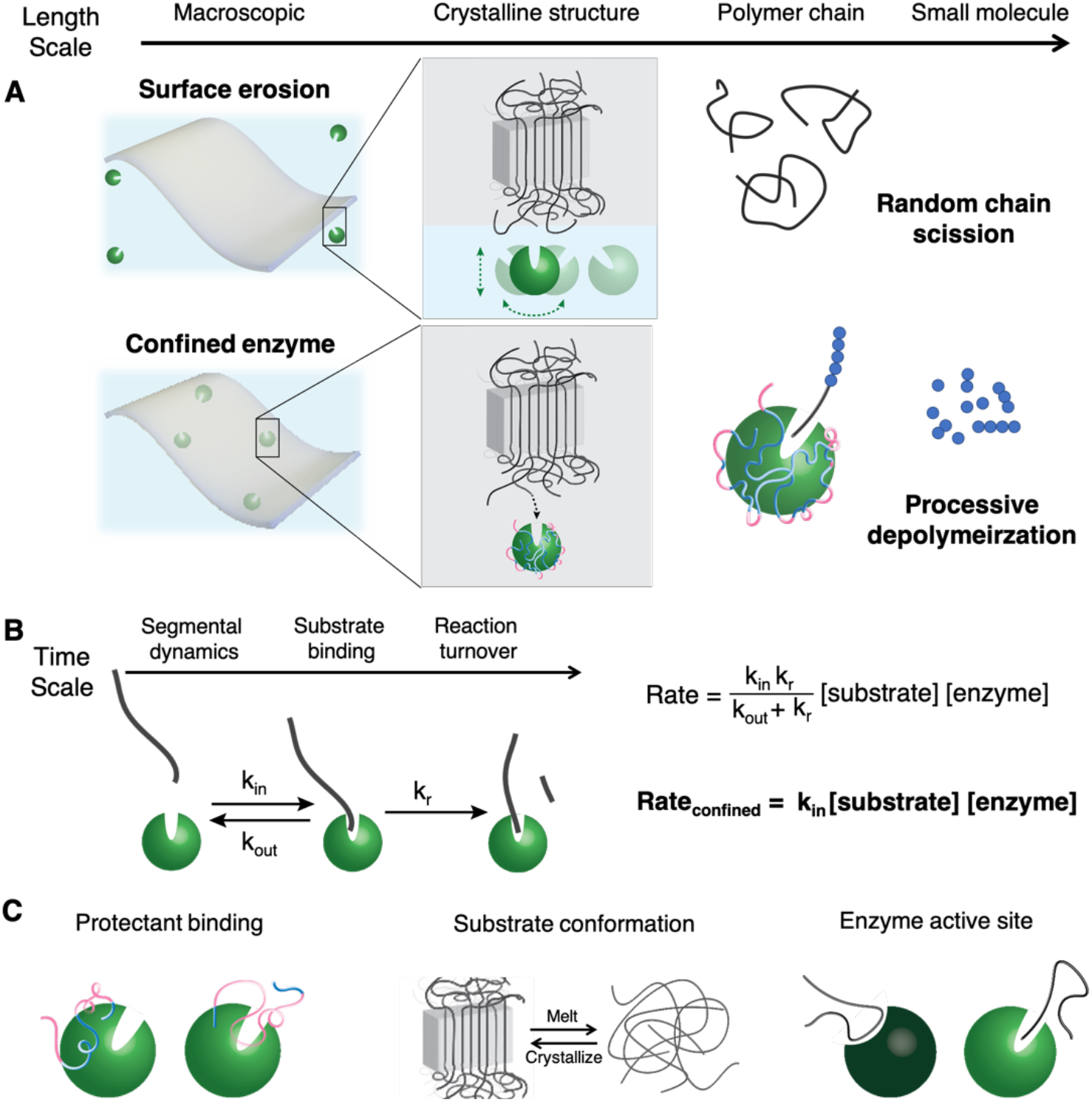
Consideration of enzymology of embedded enzyme with macromolecular substrates. (**A**) Schematically shows variation in the enzyme mobility may change the pathway of polymer degradation from random chain scission to processive depolymerization with nanoscopic confinements. (**B)** describes the reaction kinetic changes where macromolecular substrate binding becomes rate limiting factor for confined enzyme. (**C**) schematically shows additional factors to be considered to modulate enzymatic reactions toward polymer degradation.

Here, we show that distributing enzymes as nanoscopic clusters within semicrystalline polymers can realize chain-end mediated, processive depolymerization of the whole chain with repolymerizable small molecule byproducts, while preventing enzyme leaching to ensure continuous depolymerization post microplastic formation. Specifically, rapid (hours) degradation of polyesters, i.e. poly(caprolactone) (PCL) and PLA, can be realized by adding trace enzymes (~0.02 wt% lipase for PCL and 2 wt% Proteinase K for PLA). Enzyme depolymerizes the host polymer only after binding to a polymer chain end. Thermodynamically, the polymer conformational entropy and tunable segmental cooperativity enable programmable degradation without compromising integrity during processing and long-term storage. In addition to providing design insights for biocatalysis in the solid state, present studies lead to scalable bioactive plastics for efficient sorting, such as recovery of precious metal filler from flexible electronics.

A random heteropolymer (RHP) approach can effectively stabilize and solubilize enzymes in organic solvents and was used to achieve nanoscopic enzyme dispersion (*8, 10*). PCL and PLA are considered as suitable alternatives to commodity plastics (*4*), but still suffer from lengthy degradation (years) in landfills or marine environments. Lipase has been used broadly to catalyze reactions (*22*), and *Burkholderia cepacia* lipase (“BC-lipase”) was used here. Using an RHP containing the MMA/2-EHMA/3-SPMA/OEGMA molar ratio of 0.50:0.20:0.05:0.25 and BC-lipase, the RHP-lipase is well-dispersed in a range of solvents, forming ~285 nm clusters in toluene (**Fig. S1**). At 0.02 *wt*% loading, the lipase is uniformly distributed throughout the as-cast PCL films as seen in the fluorescence microscopy images (**Fig. 2a)**. The nanoclustered lipase retains over 40% activity after heating at 80 °C for 5 hours. Overlaid polarized optical microscope and fluorescence microscope images (**Fig. 2b)** show that lipase is incorporated within crystalline spherulites due to its slow diffusivity in the PCL melt compared to the crystal front growth rate. Transmission electron microscopy (TEM) images show RHP-lipase clusters, ~50 nm to ~500 nm in size, are dispersed between the crystalline PCL lamellae (**Fig. 2c)**. With up to 2 *wt*% enzyme, there are only minimal changes in PCL bulk crystallinity and mechanical properties (**Fig. S2 and Fig. S3**). Small angle x-ray scattering (SAXS) profiles show similar PCL crystallization with and without lipase incorporation (**Fig. S4**).

**Fig. 2.**
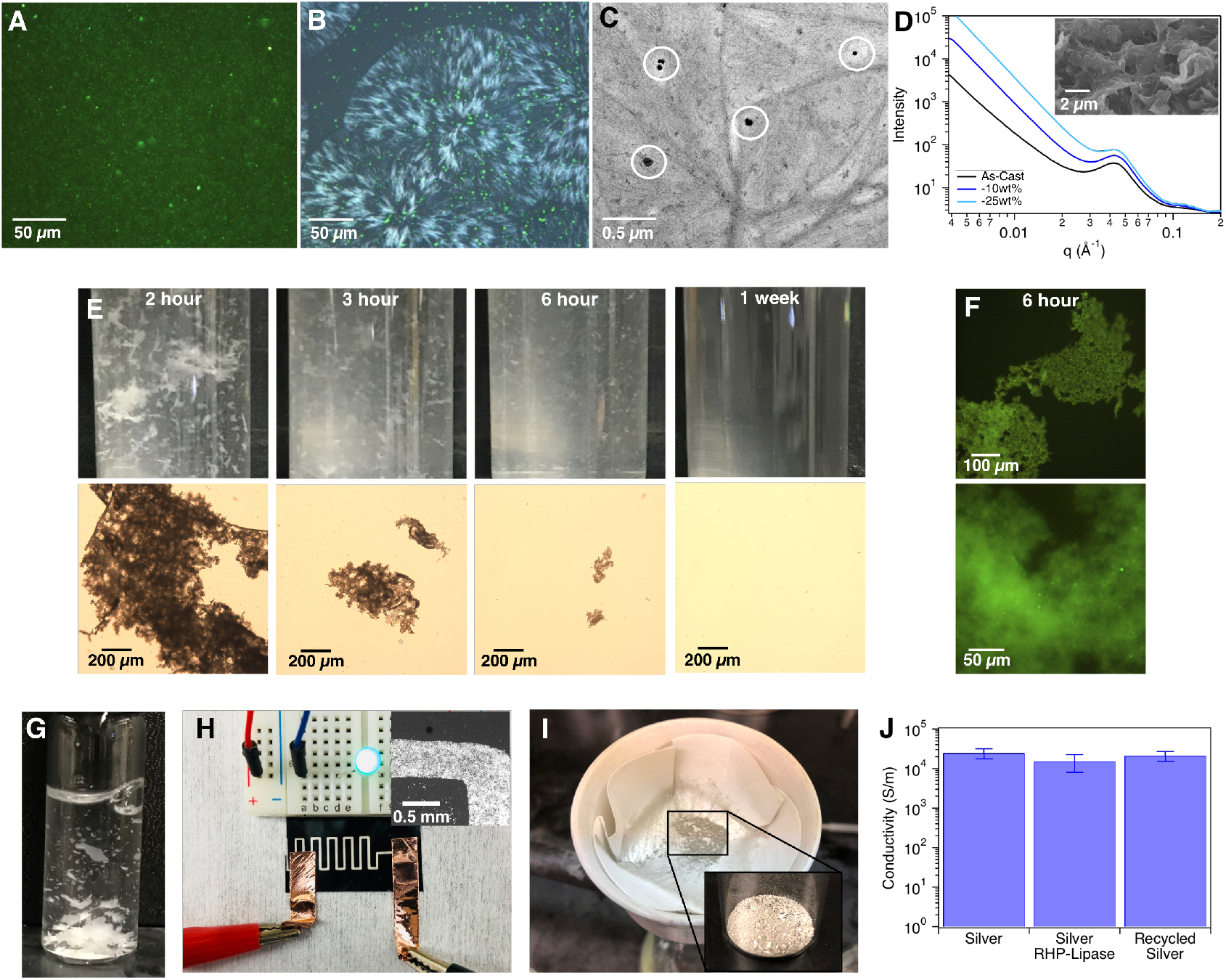
Embedding enzyme nanoclusters in PCL accelerates its degradation for convenient sorting and filler recycle. (**A**) Fluorescence microscope image showing homogeneously distributed RHP-lipase. (**B**) Overlaid fluorescence and polarized microscopy images of a melt-processed PCL-RHP-lipase film recrystallized at Tc = 49 °C. (**C**) TEM image of solvent cast PCL-RHP-lipase showing RHP-lipase clusters within semicrystalline PCL matrix. (**D**) SAXS profile confirming nanoporous structure formation during internal degradation by confined enzyme. The inset shows the crosssectional SEM image after 50% weight loss. (**E**) Photographs and optical microscope images of degradation via confined BC lipase over time showing reduction of microplastics; (**F**) fluorescence image showing confined BC lipase remains with microplastics to ensure continuous degradation; (**G**) Embedding RHP-lipase cluster in PCL/PLA blends enables sorting of PLA after selective depolymerization of PCL. (**H**) 3-D printed plastic circuit containing silver flakes and RHP-lipase with low resistance. (**I**) Silver flakes collected via simple filtration after degradation in 37 °C buffer. (**J**) Conductivity of 3-D printed plastic circuits containing just silver, silver + RHP-lipase, and recycled silver.

Semicrystalline PCL containing RHP-lipase (called “PCL-RHP-lipase”) at 0.02 *wt*% lipase loading rapidly degrades once immersed in a 40 °C buffer solution. As expected, PCL-RHP-lipase degrades internally rather than via surface erosion. **Fig. 2d** shows SAXS profiles of PCL-RHP-lipase before and after ~25% mass loss. Intensity increased in the low *q* range as degradation proceeded due to the nanoporous structure formation as internal degradation proceeds. This agrees with the cross-sectional scanning electron microscopy (SEM) image (**Fig. 2d inset**). As degradation proceeds, the PCL-RHP-lipase sample disintegrates into microplastic particles (**Fig. 2e**). Using fluorescently labeled lipase, no enzyme leaching was observed, and the lipase remains well dispersed within PCL microplastic particles and continues to degrade the host matrix (**Fig. 2f**). After 24 hours, ~95 wt% of PCL samples are degraded to small molecules and no microparticles can be observed via optical microscopy or TEM after 1 week. Control experiments using reverse micelle-encapsulated lipase at the same lipase loading stop degrading after ~40 % mass loss due to leaching, consistent with previous reports (**Fig. S5**) (*9, 17*).

The nanoscopic enzyme embedding approach is robust. Similar results were obtained using a different enzyme/polymer pair, i.e. proteinase K (*23*) and PLA, chosen as PLA is replacing polyolefins for packaging and other commodity applications (*24*). RHP-proteinase K was embedded in PLA, leading to 80 % PLA degradation to lactic acid monomer and dimer (**Fig. S6**) after 5 days in 37 °C buffer. Since lipase cannot degrade pure PLA, the enzyme selectivity was tested in polymer blends. In PCL:PLA blends (75:25 or 50:50 by weight), embedded RHP-lipase nanoclusters selectively degrade PCL after 24 hours at 37 °C for easy separation of PLA (**Fig. 2g**).

Using a commercial lipase containing protectants (lipase_cb_) without purification, RHP-lipase_cb_ forms sub ~300 nm particles in toluene (**Fig. S7**). Similar degradation results were observed once RHP-lipase_cb_ are dispersed in PCL. The scalability of confining enzyme nanoclusters in plastics enables fabrication of bioactive plastics with immediate technological relevance. PCL-RHP-lipase_cb_ was blended with silver flakes to formulate conductive ink for 3-D printing of flexible electronics (**Fig. 2h)**, an emerging class of materials (*25–27*). With nanoscopic lipase dispersion, the printed circuits have resistance less than 0.5 Ohm, sufficient to operate LEDs. After 7 months of storage at room temperature and 1 month of continuous operation with a 5V potential, the circuits degrade in 37 °C buffer, confirming the confined enzymes’ long-term stability and resistance to electrical-induced deactivation. The high purity silver flakes were collected (**Fig. 2i**) and reused repeatedly without compromising electrical conductivity (**Fig. 2j**).

PCL-RHP-lipase and PLA-RHP-protease K degradation proceeds via preferential chain-end scission rather than random chain scission found in lipase solution or reverse micelle-based dispersion. GPC analysis shows that as degradation proceeds, the PCL molecular weight remains the same with no notable formation of polymers or oligomers with reduced molecular weight (**Fig. 3a**). Liquid chromatograph-mass spectrometry (LC-MS) confirmed the primary by-products are small molecules less than 500 Da in size (**Fig. 3b and Fig. S8**). As a proof of concept, these byproducts can be readily repolymerized into PCL. In the control experiments using lipase solution (0.1 mg/mL) or reverse micelle-based lipase dispersion, a random chain scission was seen with short PCL chain by-products. To further confirm the chain end mediated depolymerization mechanism, control experiments were performed using a triblock copolymer, where the PCL is capped on both ends with a short polystyrene (PS) block (PS-PCL-PS, 1,500-8,000-1,500 g/mole). With similar treatment, PS-PCL-PS degradation was negligible after 2 days (**Fig. S9**). Thus, nanoscopically confined lipase needs to bind to PCL chain ends to initiate the depolymerization.

**Fig. 3.**
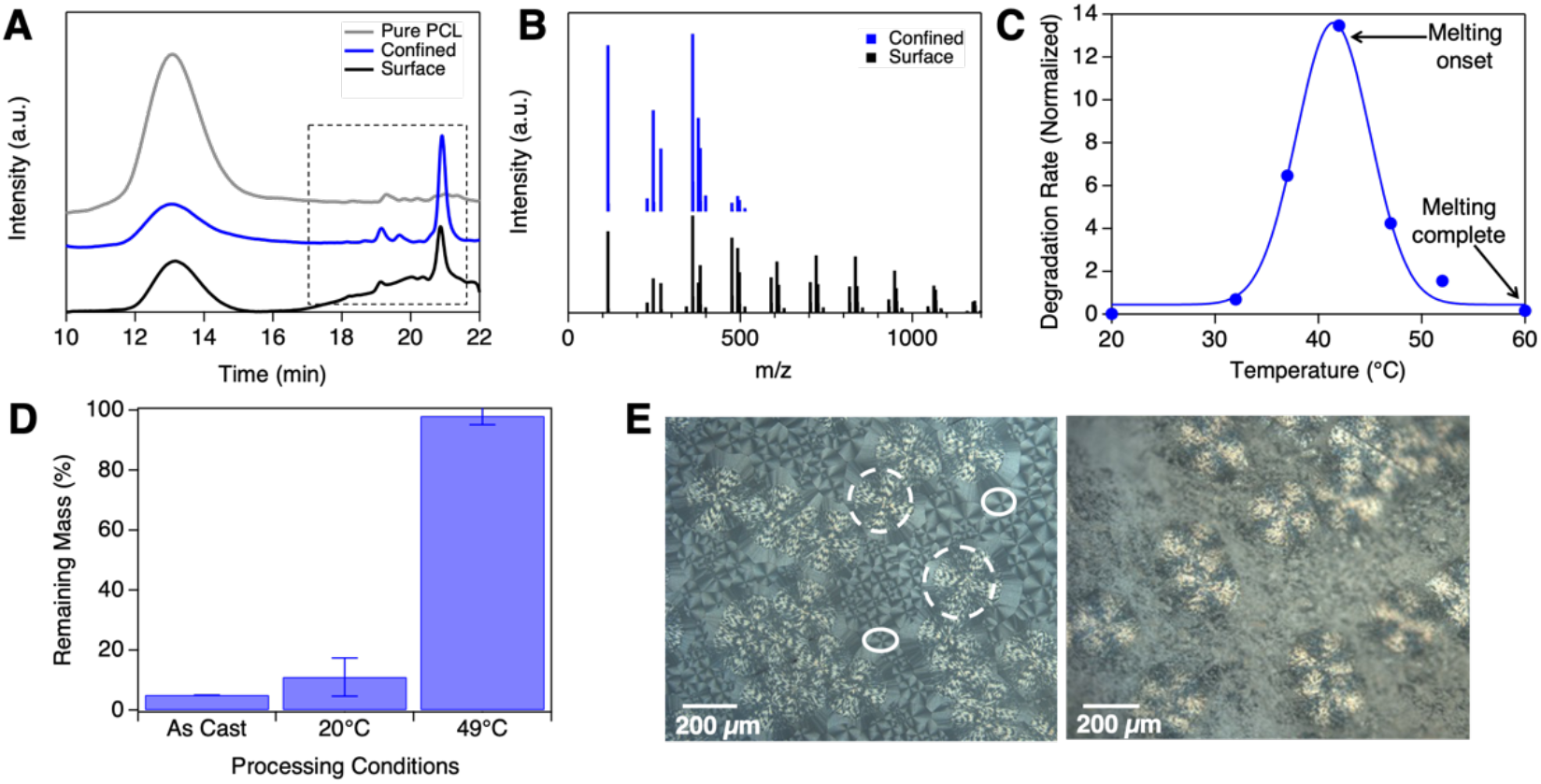
Confined lipase degrades PCL via a chain-end mediated processive depolymerization and leads to programmable degradation. (**A**) GPC of PCL after surface erosion and confined degradation that includes the remaining film and degraded by-product. (**B**) Mass spectra of surface erosion and confined degradation that include the remaining film and degraded by-product. (**C**) Confined enzyme degradation rate as a function of degradation temperature. (**D**) Remaining mass after 24 hours in 37 °C buffer for PCL-RHP-lipase films processed under different conditions. The samples crystallized at Tc = 49 °C exhibit negligible degradation up to 3 months. (**E**) (Left) Polarized optical image of a PCL-RHP-lipase film crystallized at Tc = 49 °C for 12 hours (dashed ovals) and then quenched at 20 °C for 5 minutes (solid ovals). (Right) Polarized optical microscopy image of the same film after being immersed in 37 °C buffer for 24 hours.

After embedding lipase nanoclusters, the apparent enzyme reactivity shows different temperature dependence. PCL-RHP-lipase undergoes negligible degradation at room temperature over 3 months, while lipase in solution degrades pure PCL by ~30% in just 2 days. This was attributed to the hindered mobility of the chain end due to semicrystalline morphology despite the low glass transition temperature of amorphous PCL. Degradation rate increases significantly up to the onset of PCL melting (~43 °C, **Fig. S10**) as expected from increased chain mobility. However, the degradation rate surprisingly exhibits an exponential decrease as temperature increases beyond 43 °C (**Fig. 3c**). The degradation rate diminishes when PCL is fully melted (> 60 °C). However, control experiments confirmed that the confined enzyme has a higher activity for hydrolyzing a small molecule substrate above 43 °C, which rules out enzyme denaturation (**Fig. S11**). Rather, we hypothesize that it is the global polymer conformation that affects the thermodynamic balance of chain end binding to the confined enzyme as well as the chain conformational entropy. There is a much lower conformational entropic penalty for a crystallized chain end to bind to an enzyme than an amorphous chain, provided sufficient mobility (*28*). The high entropic penalty for enzyme binding overtakes the effects of increased chain mobility, leading to large reductions in degradation rates at higher temperatures and eventually no degradation of PCL in melt state. The observed temperature dependence of apparent enzyme activity ensures polymer integrity during melt processing and during long-term storage. These observed results differ from the long-standing sentiment that crystallinity slows enzymatic degradation of both synthetic (*1, 23*) and natural polymers (*29, 30*) and could have important ramifications for the emerging field of solid-state enzymology and bioactive plastics.

Once the chain end is bound, the lipase processively catalyzes the PCL depolymerization on a single chain level. From 0 to 1 hour, the bulk percent crystallinity increases from 39 ± 1.8 % to 47 ± 3.0 % due to preferential amorphous degradation. However, as the degradation proceeds, the overall PCL crystallinity doesn’t change when the sample weight loss increases from 20% to 80% (**Fig. 4a**). Thus, the PCL segments in both the amorphous and crystalline phases are degraded as opposed to only the amorphous segments with high mobility. The results point to a single chain degradation mechanism whereby the confined lipase binds a polymer chain end and consecutively depolymerize the chain without releasing it (*i.e*. “processive degradation”). As the lipase pulls the segments in the stem spanning the crystalline lamellae, the competing force is governed by the multi-pair intermolecular interactions within the crystalline domain. Crystallization temperature (Tc) was varied to modulate the lamellar thickness—and in turn the competing force—while retaining similar bulk crystallinity and enzyme distributions (**Fig. S12 and Fig. S13**). Results showed that thicker crystalline lamellae (Tc = 49 °C) undergo negligible degradation over 3 months in 37 °C buffer while films with thinner crystalline lamellae degrade by up to ~95% in 24 hours (**Fig. 3d**). The processive degradation mechanism allows for both temporal and spatial control of PCL-RHP-lipase degradation. A film crystallized at 49 °C for 12 hours and then quenched at 20 °C exhibits two distinct crystalline morphologies (**Fig. 3e**). After being immersed in 37 °C buffer for 24 hours, only those regions crystallized at 20 °C exhibited degradation; the large spherulites that were grown at 49 °C retained their initial structure (**Fig. 3e**).

**Fig. 4.**
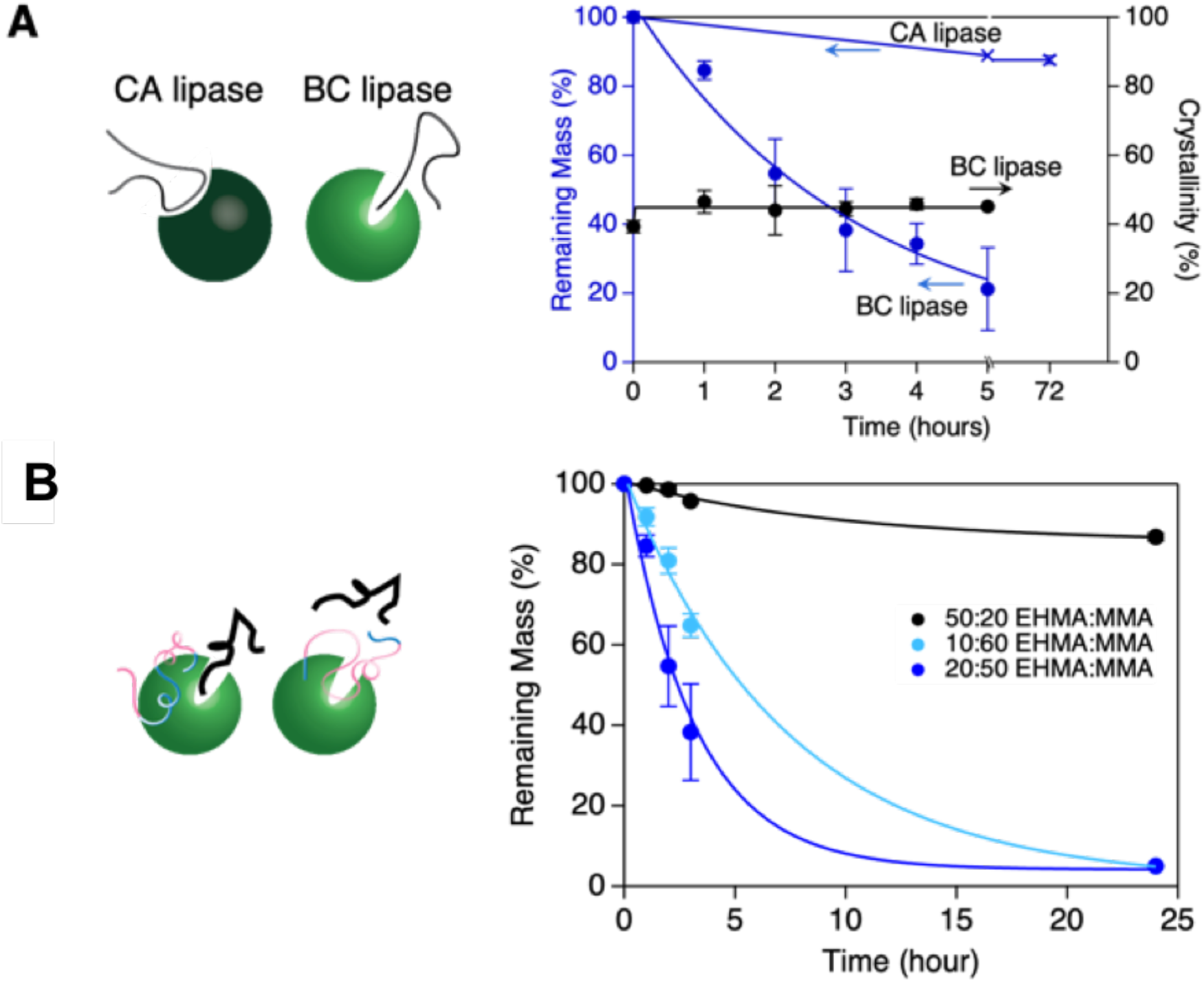
Understand design rules toward enzyme-mediated polymer degradation. (**A**) The fraction of remaining PCL as a function of degradation time for BC lipase and CA lipase, respectively. The crystallinity of remaining PCL remains similar during degradation in PCL/RHP/BC lipase blends. Only ~15% of PCL degradation was observed after 72 hrs when nanoclusters of RHP/CA lipase was embedded in PCL. (**B**) Effect of RHP composition on the apparent enzyme activity. With higher hydrophobicity of the RHP by increasing the EHMA fraction, the PCL degradation slows down. This is attributed to the strong interactions between RHP and hydrophobic surfaces around the BC lipase active site that block the polymer substrate binding as schematically shown on the left.

The BC-lipase has common traits of processive enzymes, which typically possess hydrophobic binding interactions and funnel-like active sites that facilitate single-chain sliding while hindering substrate dissociation (*29, 31*). Active site analysis shows a deep (up to 2 nm) and narrow (4.5 Å at the base) funnel from lipase’s surface to the catalytic triad (**Fig. S14a**) (*32*). This geometric restriction could exclude bulky substrates and prefer only mobile PCL chain end segments to reach the catalytic residue, which would be especially pertinent for confined enzymes that cannot easily alter their conformation to facilitate substrate binding. To test the hypothesis, a different lipase (*Candida Antarctica* lipase, “CA lipase”) was used. CA lipase has a surface-exposed active site that is quite shallow, approximately half as deep as BC lipase’s (~1nm vs. 2 nm from the surface). (**Fig. S14b**). It is expected that exposed PCL middle chain segments can be bound to the shallow binding site of the CA lipase rather than exclusively chain end binding. With the same treatment, PCL-RHP-CA lipase degradation proceeded via random scission (**Fig. S15**). The CA lipase-catalyzed degradation stops with increased PCL crystallinity after just ~12% mass loss since the lack of single chain processivity prevents crystalline stem degradation, while BC lipase processivity enables degradation of both amorphous and crystalline segments (**Fig. 4a**). Moreover, confined CA lipase has no crystalline lamellae thickness dependence nor the thermal treatment history dependence (**Fig. S16**). The difference in the apparent enzyme activity for confined BC and CA lipase demonstrates the importance of enzymes’ specific substrate interactions and reaction mechanisms for bioactive plastic design.

To achieve nanoscopic dispersion of bioactive elements in synthetic polymers, RHPs are needed for stabilization and compatibilization and are expected to be surrounding the enzyme surface. For RHP-based blends, small molecule substrates can easily penetrate the RHP protective layer to reach the active site; however, large macromolecule substrate binding requires more local movement and balancing interactions between enzyme-protectant and enzyme-substrate. The entrance to and lining of BC lipase’s active site comprise large hydrophobic regions that is important for the PCL binding. All prior results with lipase were based on the RHP design with MMA/2-EHMA/3-SPMA/OEGMA molar ratio of 0.50:0.20:0.05:0.25. Two additional RHPs were used to test the binding competition hypothesis where the EHMA to MMA ratio was varied. When the monomer ratio was changed to MMA/2-EHMA/3-SPMA/OEGMA molar ratio of 0.20:0.50:0.05:0.25, the overall hydrophobicity and that of the hydrophobic block increases. This leads to stronger interactions between the RHP and the hydrophobic surfaces of the lipase. Under the same treatment, the degradation is significantly slowed for the RHP with 50:20 EHMA:MMA, while less hydrophobic RHP compositions still enable rapid degradation (**Fig. 4b**). Thus, the enzyme-protectant interactions need to be robust enough to disperse and protect enzymes during processing but soft enough to not outcompete substrate binding interactions at the active site. Such a design rule is useful to guide enzyme-directed evolution to expand the toolbox for plastic modification or degradation (*6*).

Enzyme behavior in a solid matrix varies significantly once nanoscopically confined. Understanding enzymes in plastics not only gives new insights into solid-state enzymology with a macromolecular substrate but also enables fabrication of functional plastics with programmable life cycles. While developing new degradable plastics and green materials has great merit, considering recent developments in synthetic biology and genome information, synergizing bioactive components and synthetic polymers may open new paths to encode information toward new classes of smart materials, like nature has been doing all along.

## Funding

This work was supported by the U.S. Department of Defense, Army Research Office under contract W911NF-13-1-0232 (RHP-lipase design), the U.S. Department of Energy, Office of Science, Office of Basic Energy Science, Materials Sciences and Engineering Division under contract DE-AC02-05-CH11231 (structural characterization of PCL-RHP-lipase and 3-D printing of recyclable conductive ink) and the Bakar Fellowship (Lipase_ab_-based studies). C.D. was supported by the National Defense Science and Engineering Graduate (NDSEG) Fellowship from the U.S. Department of Defense. Scattering studies at the Advanced Light Source were supported by the U.S. Department of Energy, Office of Science, Office of Basic Energy Science under contract DE-AC02-05CH11231.

## Author contributions

T.X. conceived the idea and guided the project. C.D., J. K., P.J., L.M. and A.H performed enzyme/polymer blend studies with assistance of K.Z. and T.L. J.K and R.O.R analyzed mechanical properties. **Competing interests:** TX, CD, and JK filed provisional patent application.

## Data and materials availability

All data is available in the main text or the supplementary materials, and any additional requests can be made to the corresponding author.

## Supplementary Materials

Materials and Methods

Figures S1-S17

## Notes

### Competing Interest Statement

Ting Xu, Christopher DelRe, and Junpyo Kwon have filed a provisional patent application.

