## Supplemental Information for "Embedded Enzyme Nanoclusters Depolymerize Polyesters via Chain-End Mediated Processive Degradation"

**This PDF file includes:**

Materials and Methods

Supplementary Text

Figs. S1 to S16

### MATERIALS AND METHODS

#### Section S1. Random Heteropolymer-Lipase Embedded in PCL

#### S1.1 Enzymes

Amano PS Lipase from Burkholderia cepacia (BC lipase), Proteinase K from Tritirachium album, and Candida Antarctica Lipase B (CA Lipase) were purchased from Sigma Aldrich. Since there are surfactants solutes in the as-purchased lipase solution, the solution was purified following established procedure described in ref *(33)*. The concentration of the purified lipase stock solution was determined by using UV-vis spectroscopy based on the absorbance at 280 nm. When the enzymes were used as purchased, they are stated in the main text.

#### S1.2 Random heteropolymer (RHP)

The random heteropolymer (~70,000 g/mole with PDI= 1.3) was synthesized as previously reported *(4)*. The monomer molar composition used, unless otherwise specified (as in **Fig. 4c**), was 50% methyl methacrylate (MMA), 20% 2-ethylhexyl methacrylate (EHMA), 25% oligo(ethylene glycol methyl ether methacrylate) (OEGMA; $M_n$ = 500 g/mole), and 5% 3-sulfopropyl methacrylate potassium salt (SPMA). The RHP is referred as MMA:EHMA:OEGMA:SPMA=0.5:0.2:0.25:0.5). Two RHP variants were also used to perform control experiments and the composition is MMA:EHMA:OEGMA:SPMA=0.6:0.1:0.25:0.5 and MMA:EHMA:OEGMA:SPMA=0.2:0.5:0.25:0.5, respectively.

#### S1.3 RHP-enzyme complexes

To form the RHP-enzyme complexes, RHP and purified lipase were mixed in aqueous solution for 5 minutes at room temperature. The mixture was then flash-frozen in liquid nitrogen and lyophilized overnight to remove the water via sublimation. The remaining dried RHP-enzyme mixture was resuspended directly in the specified polymer solutions. Unless otherwise specified in the text, purified BC lipase was used to carry out studies.

RHP was mixed with BC lipase in a mass ratio of 80:1, and dynamic light scattering (DLS) was used to obtain the complex's particle size after resuspending in toluene (the processing solvent used for PCL). DLS was run on a Brookhaven BI-200SM Light Scattering System using a 90° angle. The resulting particle size distribution of RHP-lipase in toluene is shown in **Fig. S1**.

### S1.4 Processing and characterizing enzyme-embedded PCL via solution casting

PCL (80,000 g/mole, PDI < 2) was purchased from Sigma Aldrich and used without further purification. PCL was dissolved in toluene at 4wt% concentration and stirred at 55°C for at least 4 hours to ensure complete dissolution. The polymer solution was then cooled to room temperature before resuspending the dried RHP-lipase complexes directly in the polymer solution at the specified enzyme concentration. Mixtures were vortexed for ~5 minutes before being cast directly on a glass plate and air dried in a chemical fume hood.

To probe the bulk distribution of enzyme in the films, purified lipase was fluorescently labeled. Commercially-available NHS-Fluorescein (5/6-carboxyfluorescein succinimidyl ester) was used to label lipase by following the commercial procedure. The solution was centrifuged in a 15 mL 10,000 g/mole molecular weight cutoff filter for at least 3 cycles to remove excess dye from the labeled lipase. A U-MWBS3 mirror unit with 460-490 nm excitation wavelengths was used to take the fluorescence microscopy images, as shown in **Fig. 2a**. For nanoscale characterization of the complexes within the film, TEM images were taken on a JEOL 1200 microscope at 120 kV accelerating voltage. 5 *wt*% ruthenium tetroxide solution was used to stain the RHP-lipase and the amorphous PCL domains, displayed in **Fig. 2c**.

To ensure the bulk properties of the film were not compromised after enzyme incorporation, crystallinity and mechanical properties were probed via differential scanning calorimetry (DSC) and tensile testing, respectively. For DSC, ~5 mg films were pressed into aluminum pans and heated from 25 °C to 70 °C at a 2 °C $min^{-1}$ scan rate. To quantify percent crystallinity, the sample's enthalpy of melting was divided by 151.7 J $g^{-1}$, enthalpy of melting for 100% crystalline PCL *(35)*. As shown in **Fig. S2,** there is no noticeable changes in the melting temperature and the percent crystallinity after incorporating up to 2 wt% enzyme. For uniaxial tensile tests, PCL solutions were cast directly in custom-designed Teflon molds with standard dog-bone shapes. As shown in **Fig. S3,** there is <10% reduction in the modulus of PCL after embedding 2wt% enzyme. Finally, for small angle x-ray scattering (SAXS) studies, ~300μm thick films were cast in Teflon beakers. Samples were vacuum dried after degradation for at least 16 hours prior to running SAXS, which was conducted at beamline 7.3.3 at the Advanced Light Source (ALS) at the Lawrence Berkeley National Laboratory. X-rays with 1.24 Å wavelength and 2s exposure times were used. The scattered X-ray intensity distribution was detected using a high-speed Pilatus 2M detector. Images were plotted as intensity (I) vs q, where $q = (4\pi/\lambda) \sin(\theta)$, $\lambda$ is the wavelength of the incident X-ray beam, and $2\theta$ is the scattering angle. The sector-average profiles of SAXS patterns were extracted using Igor Pro with the Nika package. **Fig. S4** shows a similar long period for semicrystalline PCL with and without RHP-lipase. The same SAXS method was used to analyze the nanoporous structure of samples at different time points in their degradation, as shown in **Fig. 2d**. To obtain the cross-sectional SEM image shown in the **inset** to **Fig. 2d**, the degrading film was rinsed and fractured in liquid nitrogen to minimize fracture-induced morphological changes. The film was then mounted on an SEM stub and sputter coated with platinum. A Hitachi S-5000 SEM was used to obtain images of the film's cross section after degradation.

### S1.5 Degradation of PCL-RHP-lipase and small molecule surfactant / surface erosion control

Degradation was carried out in sodium phosphate buffer (25 mM, pH 7.2). The mass loss for timepoints up to 5 hours was determined by drying the remaining film and measuring mass on a balance after vacuum drying. The mass loss at 24 hours could not be measured on a balance because there was so little mass remaining and the remaining plastic was too small to isolate; thus, mass loss at 24 hours was estimated by lyophilizing the entire contents of the vial, running gel permeation chromatography (GPC; described in detail in section S3.1), manually integrating GPC peaks, and dividing the main remaining peak area by the area of as-cast films.

The microplastic experiment shown in **Fig. 2e** was run with a 10 mg PCL-RHP-lipase film (0.02 wt% enzyme) in 3 mL of buffer at 40 °C. The same experiment was run with fluorescently-labeled enzyme to track the enzyme in the film / microparticles during degradation (**Fig. 2f**). The same experiment was also run in 1 L of buffer while shaking the bottle periodically every few hours—the film degraded at the same rate (- 95% over 24 hours) at 1 L and 1 mL. This confirms that PCL-RHP-lipase degradation has no reliance on buffer volume, consistent with primarily internal degradation and limited enzyme leaching (**Fig. S5**).

As a control to reproduce experiments detailed in previous literature *(9, 17)*, Tween 80 was mixed with purified lipase in a 1:1 mass ratio. The resulting films were cast on glass slides, and degradation was carried out in 1 L buffer to probe the effects of leaching. In 1 L buffer, films with small molecule-embedded enzyme at the same enzyme loading as PCL-RHP-lipase degraded by ~40% in 1 day and then stopped degrading, whereas in 1 mL buffer the small molecule-embedded film degraded similarly as RHP-embedded film (~95% in 24 hours). This reliance on buffer volume shows that small molecule surfactant-embedded enzyme experiments previously reported in literature exhibit significant leaching, and in large volumes this enzyme leaching prevents complete degradation of the film.

As a further control, pure PCL films were placed in 1 L buffer with an equivalent mass of total lipase as was present in the 10 mg PCL-RHP-lipase films. Pure PCL films exhibited negligible degradation in 1 L buffer over a week, whereas pure PCL films in 1 mL buffer with the same enzyme mass lost ~80% mass in 1 day. This buffer volume dependence is expected, because enzyme must diffuse to plastic surface in order to hydrolyze the plastic.

### Section S2. Embedding Commercial Enzyme Blends in Plastics

BC lipase does not need to be purified to be resuspended in organic solvents and embedded in plastics as nanoclusters. The commercial blend, when mixed with RHP in a 2.5:1 mass ratio and lyophilized in the same procedure as the purified enzyme, forms sub-300 nm clusters in toluene

(**Fig. S6**). The confined blend, when loaded at 2 wt% total (~0.02 wt% enzyme), degraded similarly to the confined purified enzyme.

Similarly, a commercial blend of proteinase K was embedded in PLA using a 4wt% PLA in dichloromethane solution (Poly(L-lactide), 85,000-160,000 g/mole, Sigma Aldrich). When this PLA film is placed in 37 °C buffer, it degrades by ~80% over 5 days and produces primarily monomer and dimer as the by-product, as confirmed by NMR (**Fig. S7**). These NMR spectra were acquired on Bruker NEO-500 instrument (500 MHz). Chemical shifts were reported relative to the solvent peak (D2O = 4.79 ppm for 1H).

The commercial blends can be used to create relevant materials, like the flexible 3D printed electronic shown in **Fig. 2h**. PCL-RHP-lipase-silver ink was printed at room temperature from a PCL/toluene (20 *wt*%) solution containing RHP-lipase and silver flakes. Conductivity was measured using a homemade direct current setup.

### Section S3. Polymer Degradation via Confined Purified Lipase (PCL-RHP-lipase)

#### S3.1 By-product analysis

The purified lipase confined in PCL via RHP displays a processive mechanism. Gel permeation chromatography (GPC) measurements were obtained using a total concentration of 2 mg/mL of remaining film and by-product in THF. 2 uL of solution was injected into an Agilent PolyPore 7.5x300 mm column. Liquid chromatography-mass spectrometry (LC-MS) measurements were obtained by resuspending degradation supernatant in acetonitrile/water (67/33 vol%) and running through an Agilent InfinityLab EC-C18, 2.7 µm column. The mass spectrum shown in **Fig. 3b** is a combination of the major peaks seen in the chromatogram (**Fig. S8**). Degradation products were dried via lyophilization overnight before resuspending in the proper solvent for GPC or LCMS. The by-products were repolymerized using a previously-reported method *(34)* after recovering degraded PCL by-product from enzyme and buffer salts via phase extraction and filtration. The control experiment with PS-b-PCL-b-PS (cast from a solution of 15wt% polymer in toluene) was run the same way as the pure PCL experiments, and GPC (**Fig. S9**) shows negligible degradation even after 2 days at 37 °C buffer.

#### S3.2 Temperature dependence

Ramping temperature from room temperature up to ~43°C results in increased degradation rates (**Fig. S10**). Further increases in temperature, however, result in degradation rate decreases. To rule out enzyme denaturation as the cause of this degradation rate decrease, an assay was employed with a small molecule ester. PCL films containing RHP-lipase were submerged in 0.5 mM 4-nitrophenyl butyrate solution in sodium phosphate buffer at the given temperature. Activity was quantified by using UV-vis spectroscopy to monitor the absorbance over 20 minutes at 410 nm, which is where the hydrolyzed by-product absorbs. Extinction coefficient for by-product quantification was estimated as 16,500 $M^{-1}$ $cm^{-1}$. Controls of just the 0.5 mM ester solution were run at each temperature to ensure that the ester was not self-hydrolyzing. As shown in **Fig. S11**, the activity toward the small molecule significantly increases above 43 °C, ruling out denaturation as the cause for reduced PCL degradation.

#### S3.3 Crystallinity changes during degradation

At each timepoint from 0-5 hours, PCL-RHP-lipase remaining films were dried and analyzed via DSC. Despite significant mass loss (80% remaining mass to 20% remaining mass from 1-5 hours), the percent crystallinity remains constant, confirming degradation of crystalline and amorphous regions with similar rates.

#### S3.4 Melt processing to program degradation

Films were cast on microscope slides from their respective solutions. Dried films were then placed on a hot plate at 80 °C for 5 min to ensure complete melting, then crystallized at the specified temperature for up to 3 days to ensure complete recrystallization (checked using DSC and polarized optical microscope). DSC results (**Fig. S12**) indicate a similar bulk percent crystallinity, but an increase in lamellae thickness for Tc = 49 °C films (as shown by the large increase in Tm, since Tm is directly proportional to crystalline lamellae thickness based on the Gibbs-Thompson equation). Moreover, SAXS profiles show an increase in the long period for Tc = 49 °C films (**Fig. S13**).

### Section S4. Enzyme Active Site Affects Degradation by Confined Enzymes

#### S4.1 Enzyme structural analysis

Crystal structures of BC lipase (**Fig. S14A**) and CA lipase (**Fig. S14B**) are taken from entries 3LIP and 1TCA in protein data bank, respectively. A mesh depiction was used to render the structures in PyMOL. Hydrophobic residues (purple) are defined as the following amino acids: alanine, glycine, valine, leucine, isoleucine, phenylalanine, methionine, and proline. The rest were considered hydrophilic (gray).

### S4.2 Differences in confined enzyme activity: BC lipase vs. CA lipase

CA lipase blend was embedded in PCL using a 2.5:1 mass ratio of RHP:CA lipase at a similar enzyme loading to BC lipase. Degradation seemingly proceeded via completely random scission, as indicated by a shift of GPC curve to lower average molecular weight (**Fig. S15**). Moreover, films recrystallized at Tc = 49 °C degraded to the same extent as the as-cast films and show a disrupted spherulite structure after degradation (**Fig. S16**), confirming that degradation by confined CA lipase doesn't depend on lamellae thickness. Control experiments using BC lipase at RHP:BC lipase were performed and the results show no change in PCL molecular weight and there is no dependence of degradation on the crystallization temperature.

**Fig. S1.**

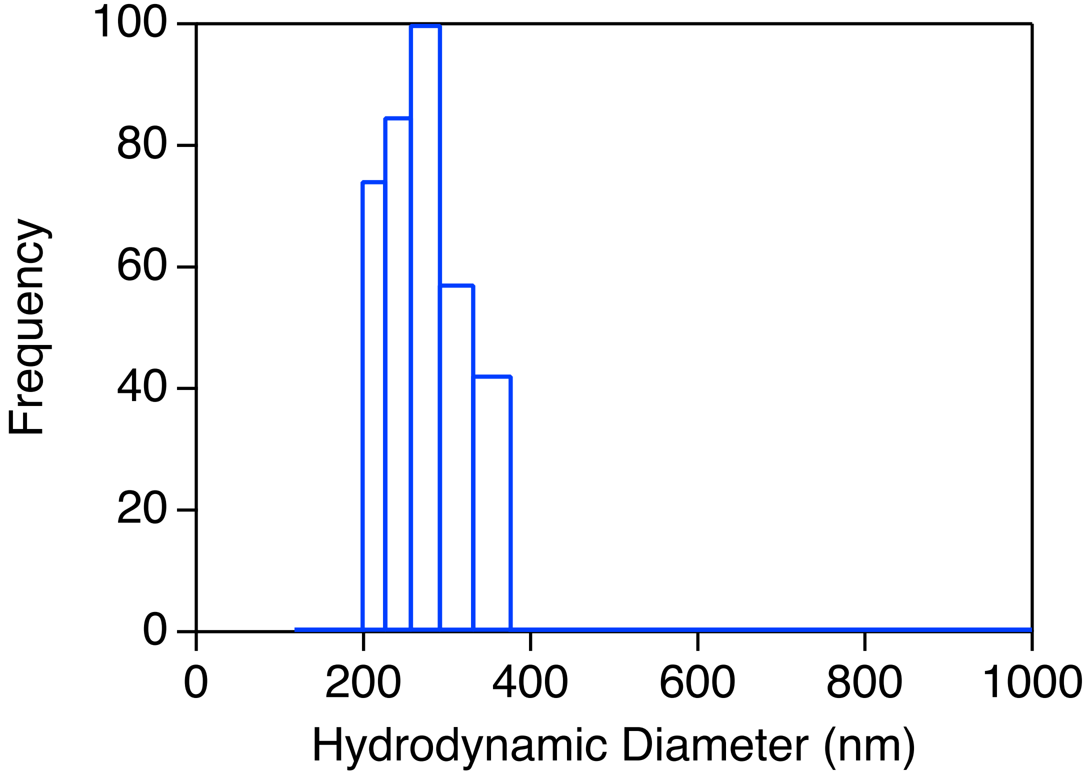


DLS of RHP-lipase in toluene with an average hydrodynamic diameter of 285 nm ± 35 nm; experiment was run at a 90° angle in filtered toluene

**Fig. S2.**

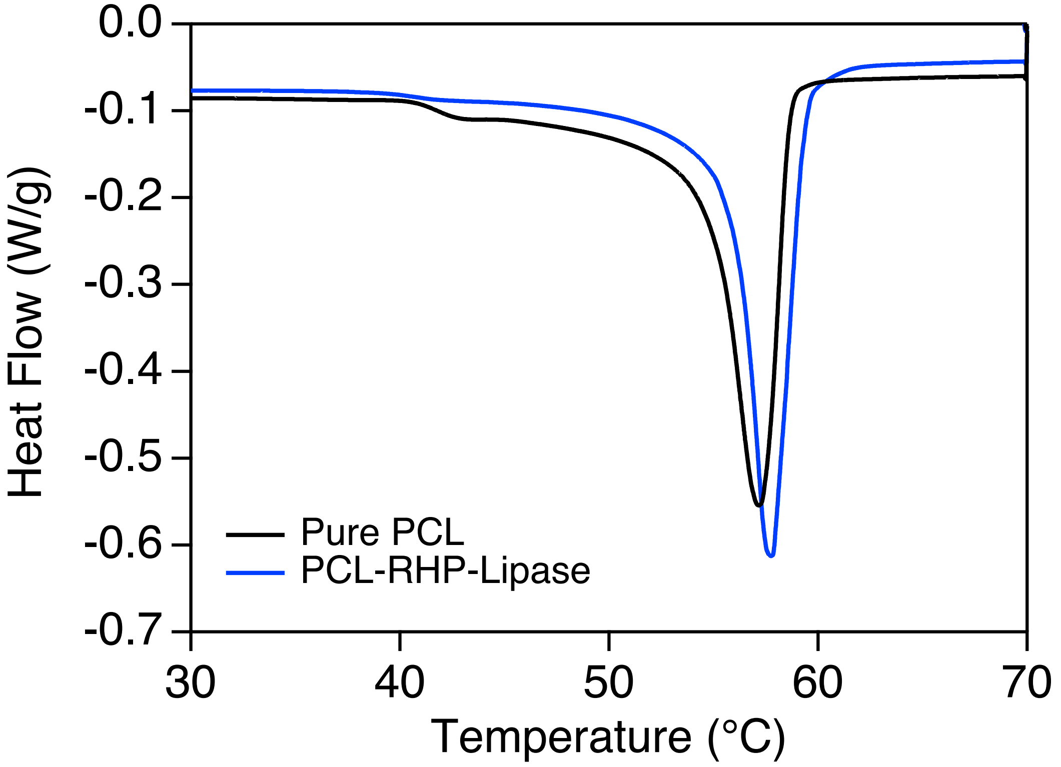


DSC curves of pure PCL and PCL-RHP-lipase as-cast films heated at 2°C/min rate

**Fig. S3.**

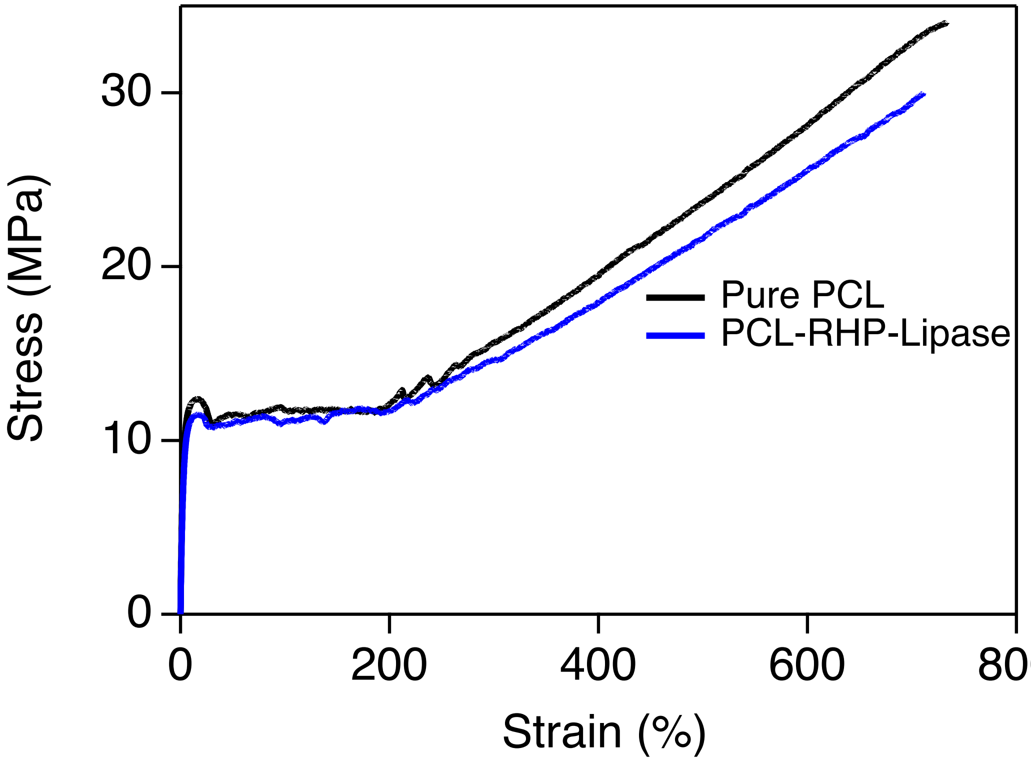


Engineering stress-strain curves from uniaxial tensile tests of pure PCL and PCL-RHP-lipase

**Fig. S4.**

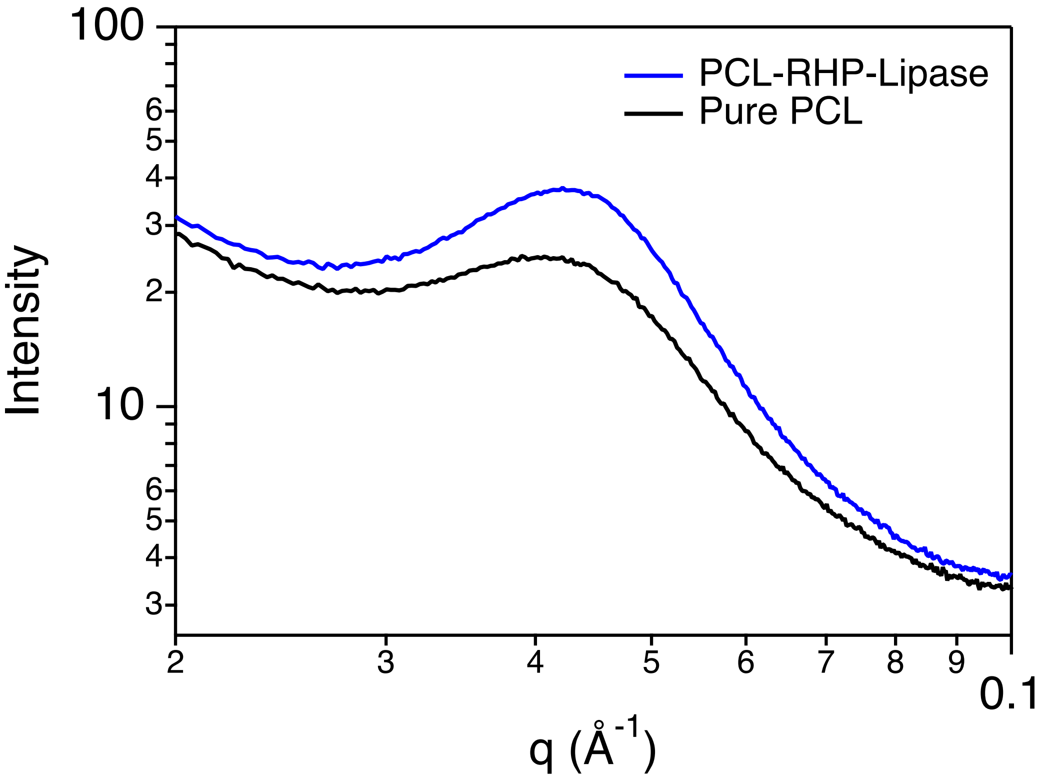


SAXS profiles of pure PCL and PCL-RHP-lipase solution cast films, run using conditions described in preceding supporting material text

Fig. S5.

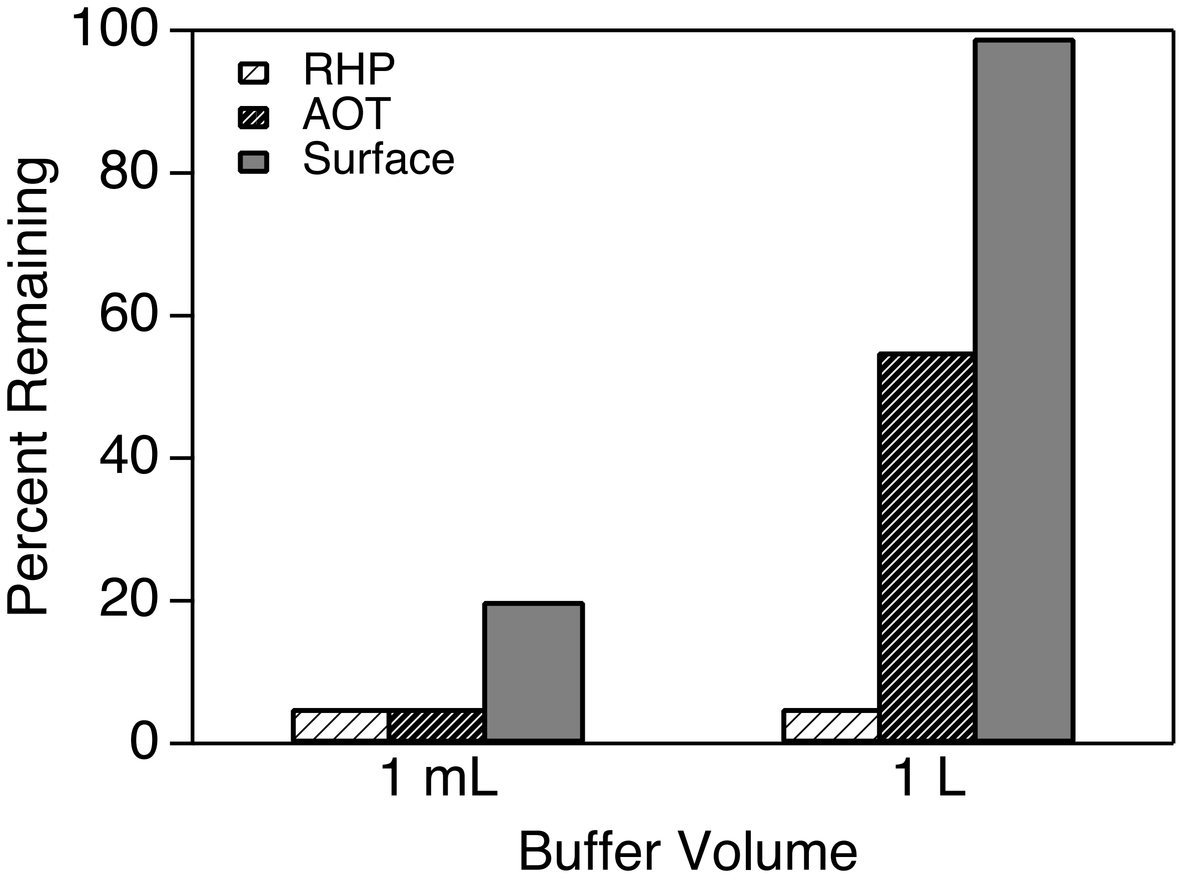


PCL degradation as a function of enzyme distribution and buffer solution volume. Embedded enzymes using either reverse micelle (AOT) or RHP can effectively accelerate PCL degradation in small buffer volume. This was attributed to the high enzyme concentration in solution that leach out form the PCL. The effect of leach out enzyme is significantly reduced when the buffer volume is increased to 1L.

**Fig. S6.**

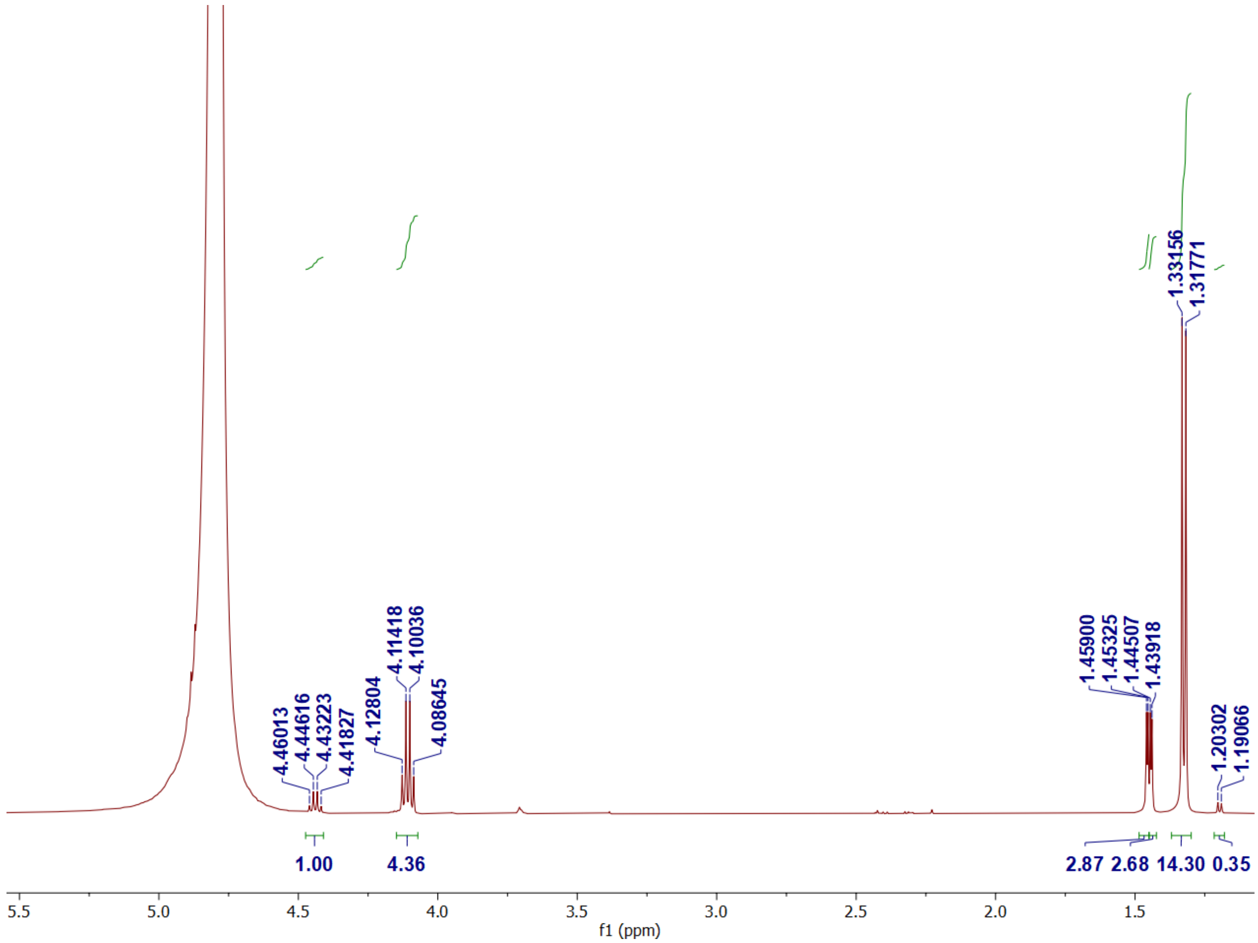


[1]H NMR run in $D_2O$ (4.79 ppm) of PLA-RHP-Proteinase K degradation by-product after 5 days in 37°C buffer demonstrating primarily monomer and dimer of lactic acid. Monomer was attributed to $\delta$ = 1.3 ppm and 4.1 ppm, $J$ = 6.93 Hz, and dimer was attributed to $\delta$ = 1.4 ppm and 4.4 ppm, J = 6.97 Hz. Monomer:dimer ratio was calculated as 84:16.

**Fig. S7.**

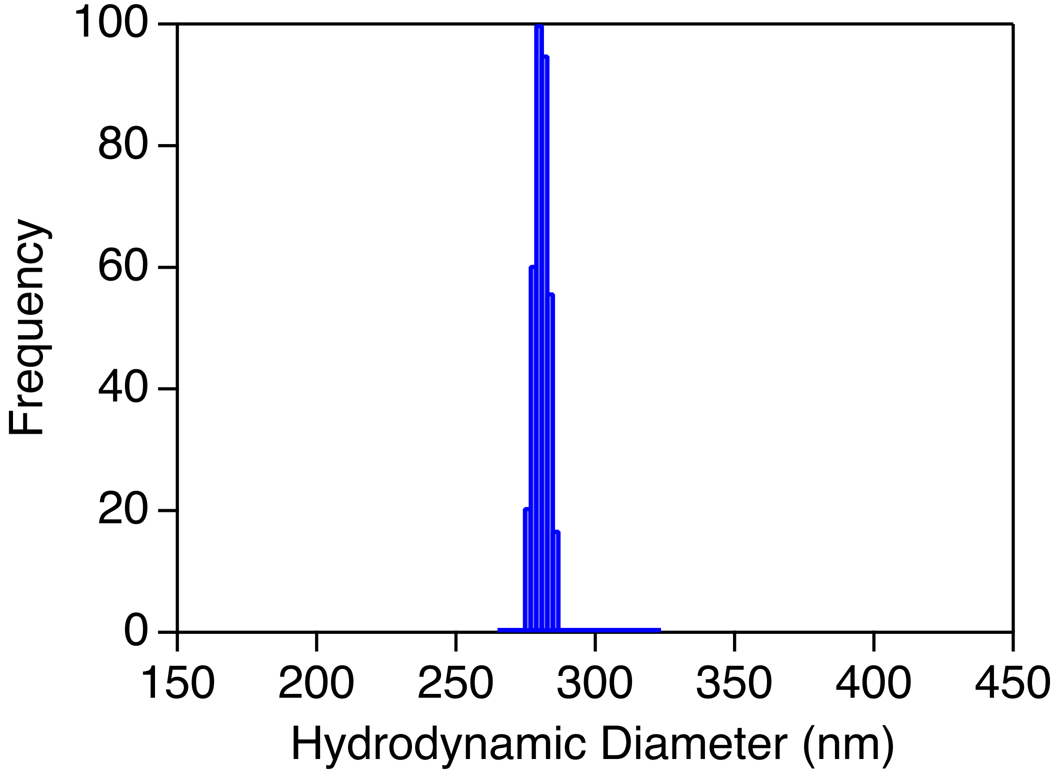


DLS of RHP-lipase$_{cb}$ in toluene showing sub-300 nm dispersion; experiment was run at a 90° angle in filtered toluene

**Fig. S8.**

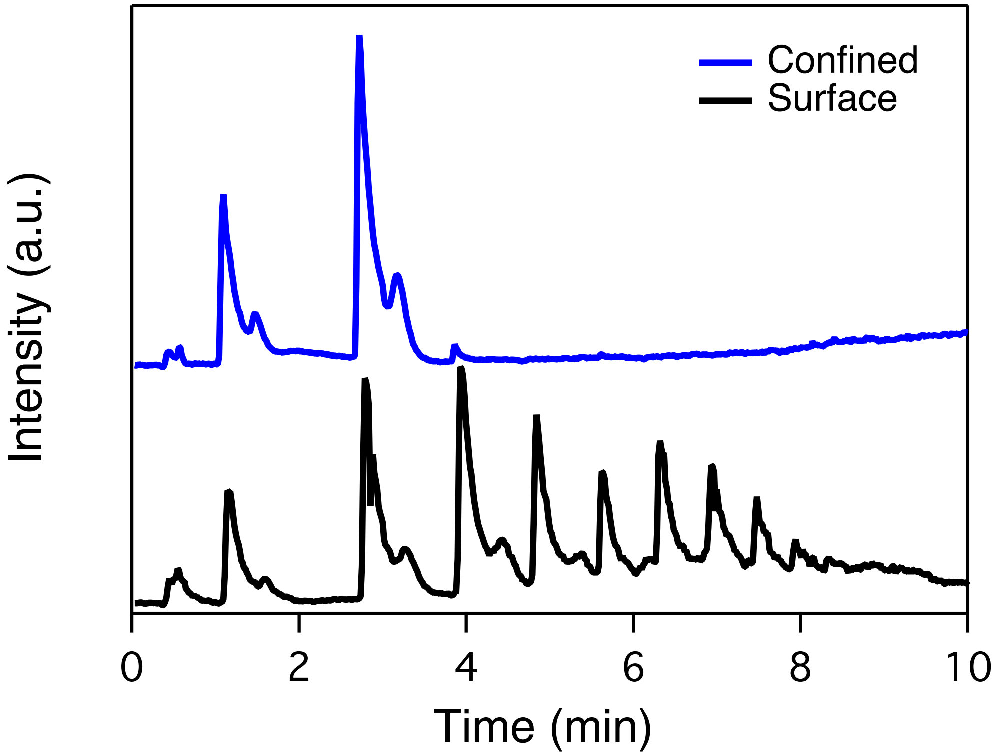


Liquid chromatogram of PCL-RHP-lipase (blue) and pure PCL in concentrated lipase blend solution (black)

**Fig. S9.**

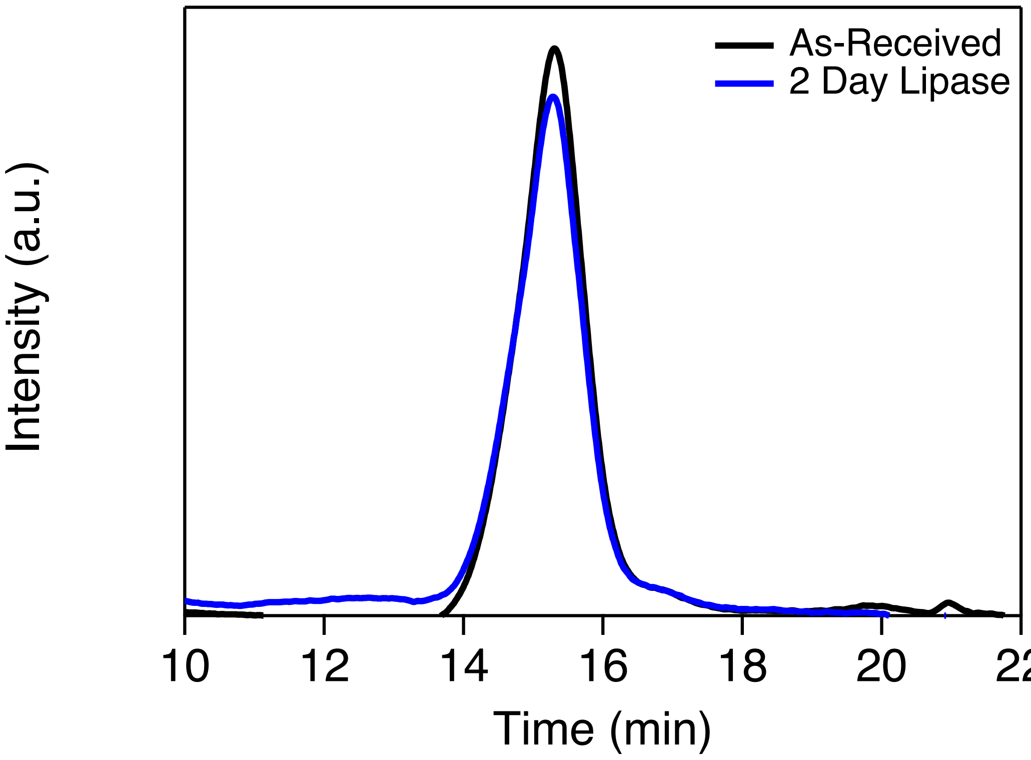


GPC of PS-PCL-PS after RHP-lipase treatment; additionally, no mass change of film after lipase treatment was detected.

**Fig. S10.**

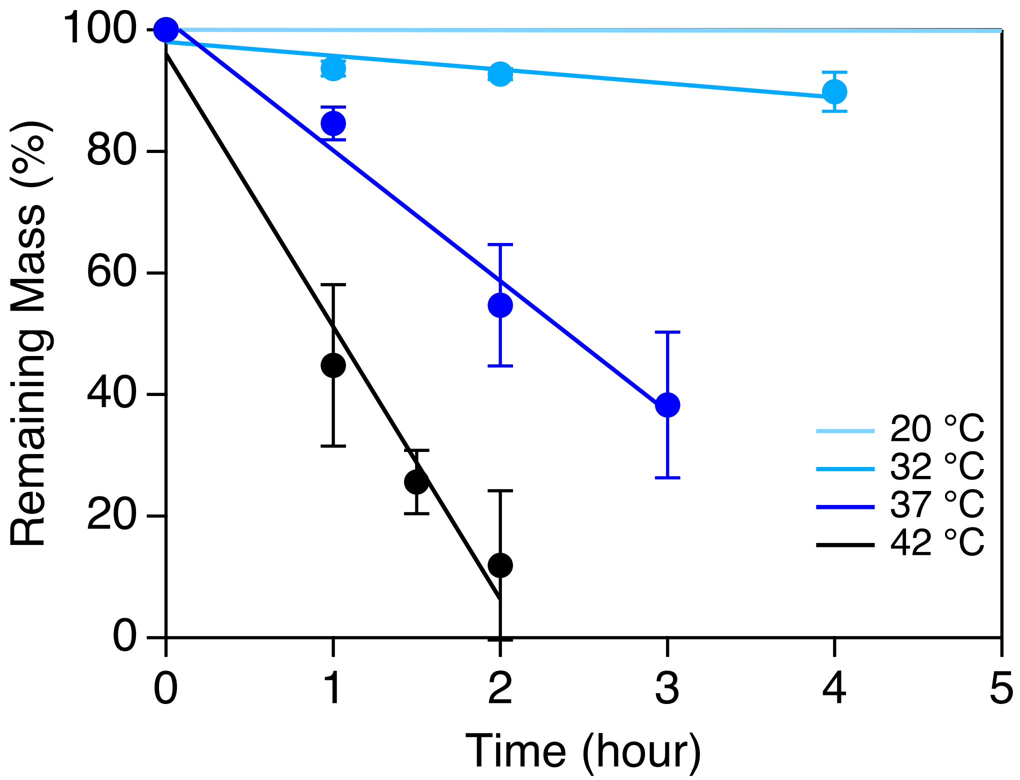


Degradation rate of PCL-RHP-Lipase as function of temperature from 20-42 °C; at each timepoint, remaining film was removed from vial, vacuum dried, and weighed on a balance

**Fig. S11.**

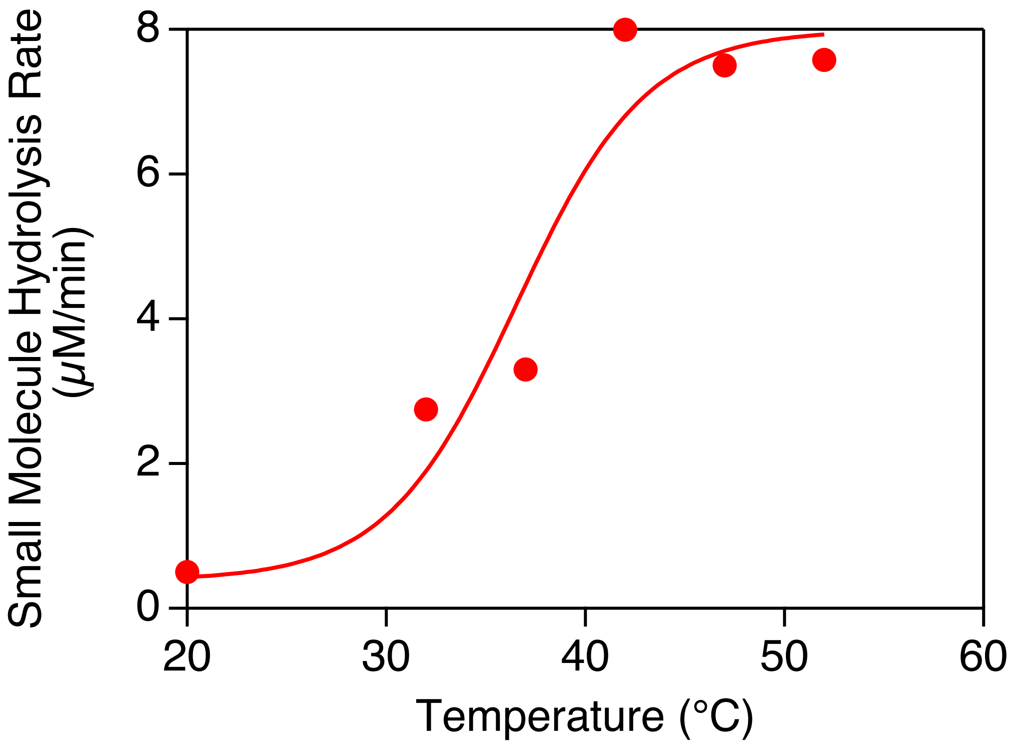


Small molecule ester hydrolysis by embedded lipase as a function of temperature; activity remained high at 60 °C but was not quantified because the film shriveled due to melting and thus was much thicker than films at any lower temperature

**Fig. S12.**

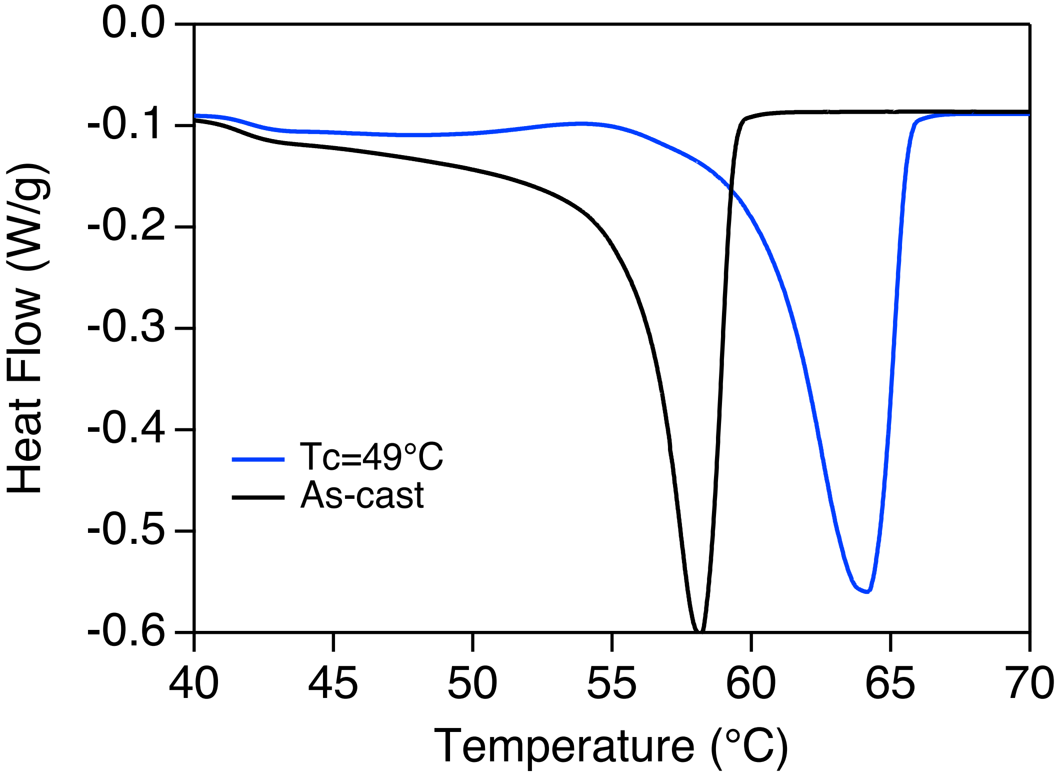


DSC curves of PCL-RHP-lipase with different recrystallization conditions run using a 2°C/min heating rate (Tc = 49 °C film has percent crystallinity of 41% ± 1.2% compared to 39% ± 1.8% for as-cast); the increase in melting temperature from ~58 °C to ~64 °C indicates a substantial thickening in crystalline lamellae for Tc = 49 °C films

**Fig. S13.**

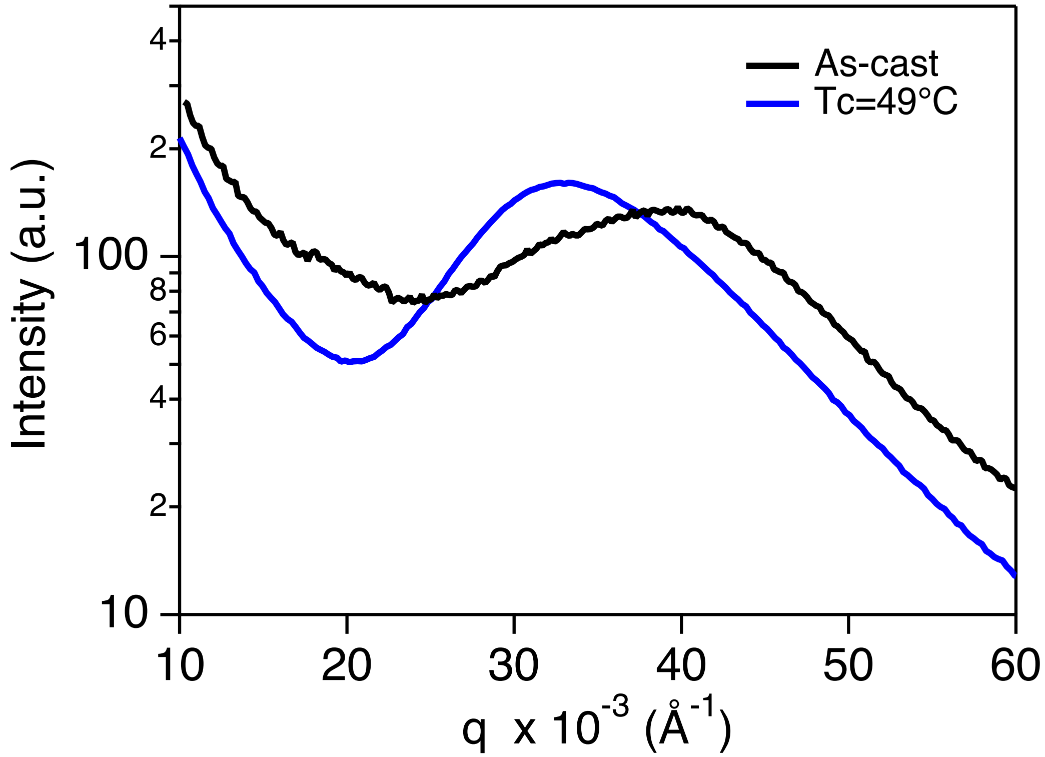


SAXS profiles of as-cast and Tc = 49°C films of PCL-RHP-lipase; the increase in long period (shift to lower $q$) combined with negligible difference in bulk percent crystallinity based on DSC data confirms a thickening in crystalline lamellae after crystallizing at Tc = 49°C

**Fig. S14.**

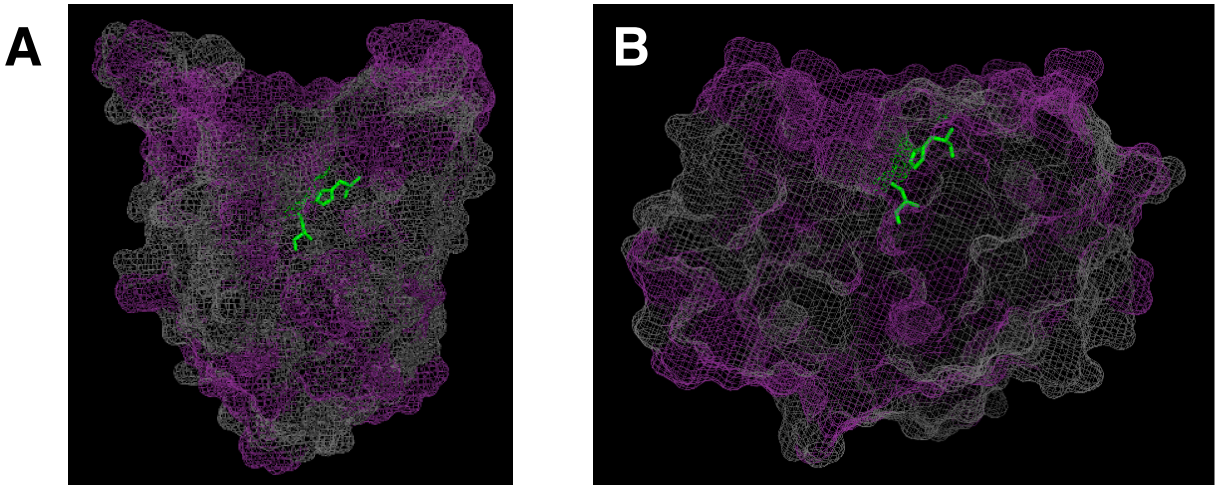


Crystal structure renderings of (A) BC lipase (3LIP on protein data bank) and (B) CA Lipase (1TCA on protein data bank). Hydrophobic protein residues are depicted in purple, while hydrophilic residues are depicted in gray. The serine and histidine residues of the catalytic triad are shown in their stick representations in green. Binding pocket for both proteins is oriented to the top of the image.

**Fig. S15.**

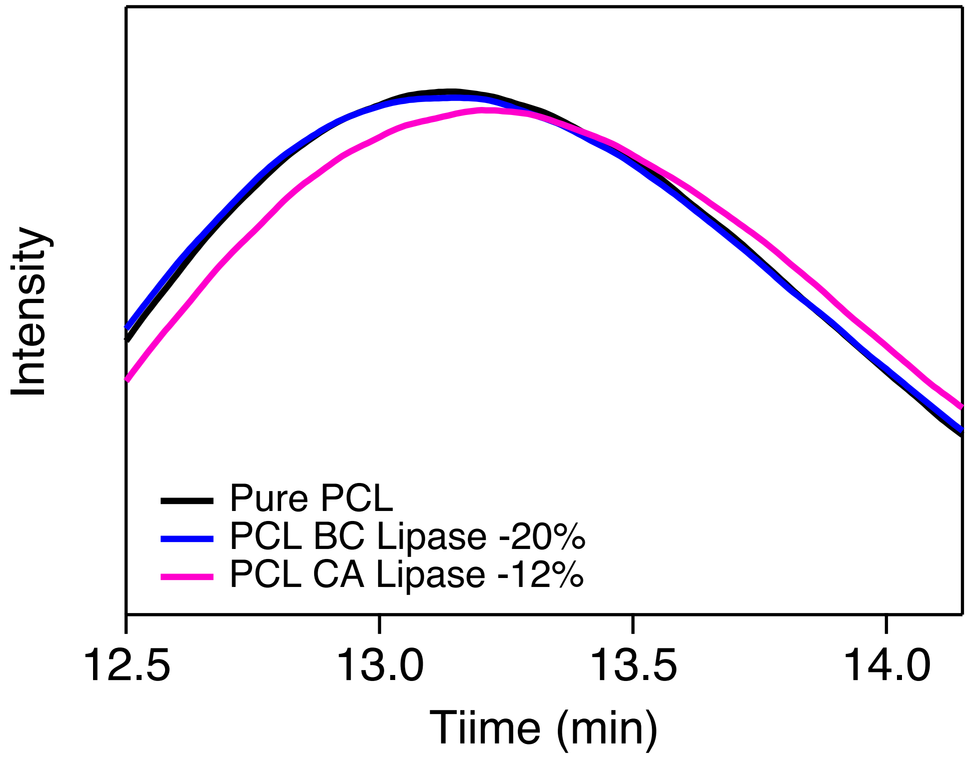


GPC curves of remaining film after specified degradation for BC and CA lipase

**Fig. S16.**

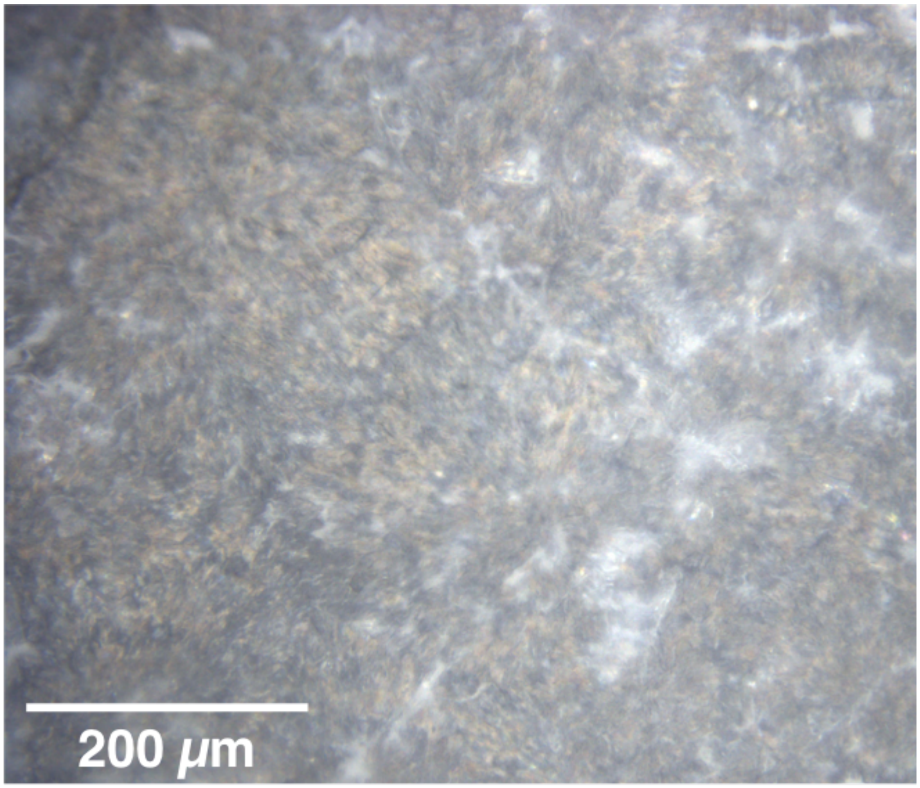


Polarized optical image of a CA lipase-embedded film crystallized at Tc = 49 °C and degraded for 24 hours in 37 °C buffer; the film degrades by the same amount as the as-cast film and the spherulites with thick lamellae are degraded, demonstrating that degradation by embedded CA lipase has no lamellae thickness or thermal treatment dependence
